# Affective Pavlovian motivation is enhanced in obesity susceptible populations; implications for incentive motivation in obesity

**DOI:** 10.1101/657833

**Authors:** Rifka C. Derman, Carrie R. Ferrario

## Abstract

Global obesity rates continue to rise, presenting a major challenge to human health. Efforts to uncover the drivers of this epidemic have highlighted the contribution of Pavlovian motivational processes to overeating. In humans, brain and behavioral reactivity to food related stimuli positively correlates with subsequent weight gain. In concordance with this, selectively bred obesity-prone rats exhibit stronger cue-triggered food-seeking via single outcome Pavlovian-to-instrumental transfer (SO PIT) than obesity-resistant rats. These data show that Pavlovian motivation is stronger in selectively bred obesity-prone groups. However, whether obesity susceptibility in outbred populations is associated with enhanced PIT is unknown. Moreover, PIT can arise via two neurobehaviorally dissociable processes, a sensory specific versus a general affective process that cannot be distinguished by SO PIT. Thus, it is unclear which PIT process is enhanced in obesity-prone groups. Therefore, we determined whether obesity susceptibility in outbred populations is associated with enhanced Sensory Specific (SS) PIT or General PIT and whether expression of these forms of PIT differs between selectively bred obesity-prone versus obesity-resistant rats. We find that in outbred rats, the magnitude of General PIT is positively correlated with subsequently determined obesity susceptibility. In selectively bred rats, the magnitude of General PIT was stronger in obesity-prone versus obesity-resistant groups. Jointly, these data show that enhanced affective Pavlovian motivation is tightly linked to obesity vulnerability, supporting a role for phenotypic differences in incentive motivation for the development of obesity. This has important implications for obesity prevention and for understanding the neurocircuitry mediating enhanced food-seeking in vulnerable individuals.

## I. INTRODUCTION

Obesity rates the world over continue to rise, presenting a major health concern (Finkelstein et al., 2009; N.C.D.R, 2016). At its most basic level, the cause of obesity is chronic over-consumption of hypercaloric diets leading to fat accumulation (Akiyama et al., 1996; Horton et al., 1995; Nascimento et al., 2008). Therefore, identifying factors that contribute to heightened food-seeking and consumption is a critical step toward understanding the etiology of obesity (Matikainen-Ankney and Kravitz, 2018; Stice et al., 2013). While many factors contribute, genetic variation renders some individuals more susceptible to diet induced weight gain. In humans, bodyweight and body mass indices are strongly influenced by genetics, with 80% of the population variance accounted for by genetic factors (Hur et al., 2008; Maes et al., 1997). Similar variation and heritability are found in outbred rat populations, with some individuals overeating and gaining more weight than others (Levin et al., 1997; Madsen et al. 2010). Behaviorally, phenotypic differences in sensitivity to motivational factors that influence food-seeking and feeding behaviors are thought to contribute to weight gain in susceptible populations (Alonso-Caraballo et al., 2018; Dagher, 2009; Ferrario, 2018; Stice et al., 2013). Therefore, identifying behaviors associated with obesity susceptibility is an important step toward understanding its neurobehavioral underpinnings.

Pavlovian associations can strongly influence food-seeking behaviors. For example, the smell of pizza can elicit desire that drives food-seeking and consumption. Numerous studies in humans have found that the degree to which Pavlovian stimuli influence neural activations in motivational circuits, and the desire for food are stronger in obese and obesity susceptible individuals (see Boswell and Kober, 2016 for metanalyses of existing human studies). Specifically, the magnitude of brain activations evoked by presentation of food related stimuli (i.e., food cues) is greater in overweight and obese versus healthy weight individuals, particularly in regions that mediate Pavlovian motivation such as the amygdala and the Nucleus Accumbens (NAc; Rothemund et al., 2007; Stoeckel et al., 2008). Moreover, the magnitude of activation in the amygdala, NAc and the ventral pallidum elicited by food cues is predictive of subsequent weight gain among healthy weight individuals (Burger and Stice, 2014; Demos et al., 2012; Yokum et al., 2014). These latter data in particular suggest that differences in brain responses to food cues precede weight gain in vulnerable populations. Consistent with this, we recently demonstrated that Pavlovian food cues exert greater motivational influence over instrumental food-seeking in selectively bred obesity-prone versus obesity-resistant rats prior to obesity (Derman and Ferrario, 2018). These data support the idea that there are phenotypic enhancements in sensitivity to the motivational properties of Pavlovian food cues in obesity susceptible populations.

The ability for Pavlovian stimuli to invigorate instrumental behaviors is a phenomenon known as Pavlovian-to-instrumental transfer (PIT), a classic measure of Pavlovian motivation (Cartoni et al., 2016). PIT can emerge via a sensory specific process or a general affective process that are neurally and behaviorally dissociable (Corbit and Balleine, 2005; Corbit and Balleine, 2011; Corbit et al., 2007). Sensory Specific PIT (SS PIT) is measured by contrasting the effects of a conditioned stimulus (CS) on two separate instrumental responses, where one response shares an outcome with the CS, but the other does not (Colwill and Motzkin, 1994). Thus, SS PIT is driven by shared sensory properties of a response-outcome (R-O) and stimulus-outcome (S-O) association (Colwill and Motzkin, 1994). One distinguishing psychological feature of SS PIT is its relative robustness; it is difficult to disrupt and emerges whether rats are tested satiated or hungry (Corbit et al., 2007). Neuronally, SS PIT relies on the basolateral amygdala (BLA) and the NAc Shell (Corbit and Balleine, 2005; Corbit and Balleine, 2011). On the other hand, General PIT arises when transfer emerges via the general affective properties shared by an R-O and S-O association. This effect can be observed when a CS enhances instrumental responding above baseline for an outcome other than that previously predicted by the CS (Balleine, 1994; Corbit and Balleine, 2005). In contrast to SS PIT, General PIT is more labile, with hunger promoting its expression and satiation reducing it (Balleine, 1994; Corbit et al., 2007). Neuronally, General PIT relies on the central amygdala (CA) and the NAc Core (Corbit and Balleine, 2005; Corbit and Balleine, 2011). Data demonstrating that obesity-prone rats exhibit stronger PIT than obesity-resistant rats used a single outcome PIT procedure, which does not distinguish between the SS and General PIT. Identifying which of these processes is enhanced in obesity susceptible populations is critical for understanding the psychological and neural processes that promote enhanced food-seeking in susceptible populations.

In the current study, we determined the degree to which SS and General PIT are enhanced in obesity susceptible versus resistant populations prior to obesity onset using two complementary rodent models. In Experiment 1, we tested outbred Sprague Dawley rats for expression of these behaviors before placing them on a moderately sugary and fatty junk-food diet in order to identify individual susceptibility to weight gain. We then performed correlational analyses between PIT behavior and bodyweight after 5 weeks on this diet. To our knowledge, this is the first study to examine whether basal expression of PIT prior to weight gain is correlated with subsequently identified obesity susceptibility in outbred rat populations. In Experiment 2, we determined whether SS and/or General PIT are enhanced in selectively bred obesity-prone versus obesity-resistant rats. This expands upon our previous work (Derman and Ferrario, 2018) by explicitly identifying which form of PIT is enhanced in obesity susceptible populations. Collectively these data provide key insights into the psychological aspects obesity vulnerability and point to specific neural circuits that may play a crucial role in this vulnerability.

## II. MATERIALS AND METHODS

### A. General Approaches

#### Subjects

Adult Sprague Dawley male rats purchased from Envigo were used for Experiment 1 (N=39). Adult Sprague Dawley male rats purchased from Envigo were used for Experiment 1 (N=39). Adult male obesity-prone (OP: N=15) and obesity-resistant rats (OR: N=15) were used for Experiment 2. Obesity-prone and obesity-resistant rats were bred at the University of Michigan using in a Poiley rotation system with 12 breeding pairs per line. These rat lines were originally developed by Barry Levin (1997). Rats were housed in groups of two or three and maintained on a reverse light-dark circadian cycle (12/12). Experiments were conducted during the dark phase of this cycle. All procedures were approved by the University of Michigan Institutional Animal Care and Use Committee. Further details for all procedures and housing can be found at: https://sites.google.com/a/umich.edu/ferrario-lab-public-protocols/.

#### General Training Procedures

Training and testing were conducted in standard Med Associates operant chambers housed within sound attenuating cabinets. Each chamber was outfitted with a recessed food cup into which 45mg pellets could be delivered via a tube attached to externally housed food hoppers. The food cup was equipped with an infrared emitter receiver unit that detected entries into the food cup. Two deflection-sensitive retractable levers flanked the food cup. Two speakers were mounted on the wall opposite to the food cup, one delivered a tone stimulus, and the other a noise stimulus. In addition, a click generator was also mounted externally on this same wall. LED red and infrared light strips were used as house lights to enable video recording of training and testing sessions via mini cameras mounted overhead (Surveilzone, CC156).

The training procedures used in Experiment 1 and 2 were identical. Prior to training, rats were food restricted to 85-90% of their *ad libitum* weights and maintained at this weight until the end of behavioral training and testing. Instrumental training, Pavlovian conditioning, and PIT testing were all adapted from Corbit and Balleine (2005; 2011). Table 1 provides a general description of response-outcome (R-O) and stimulus-outcome (S-O) relationships during training, and PIT testing; specific details of each component are given below. Rats were initially trained to retrieve food pellets from the food cups within the operant chambers in three separate sessions, using three distinctly flavored 45 mg pellets (Bioserv: Unflavored #F0021; Banana #F0059; Chocolate #F0299). Each session lasted 20 min, during which 20 pellets of one flavor were delivered on a variable time (VT) schedule of 60 sec (range, 30-90sec)

**TABLE I:** Experimental design of PIT Training. In the first phase, Instrumental Training, rats were trained to acquire 2 separate instrumental R-O associations. In the second phase, Pavlovian Conditioning, rats were conditioned with 3 separate CS-O associations, where 2 of the outcomes overlapped with the outcomes from instrumental training (O1 and O2). In the final phase, PIT Testing, rats were given access to the levers and presented with the CSs to measure the influence of CS presentation on instrumental responding. Testing was conducted under extinction conditions.

| Instrumental Training | Pavlovian Conditioning | PIT Testing |
| --- | --- | --- |
| R1-O1 | CS1-O1 | CS1: R1 v R2 |
| R2-O2 | CS2-O2 | CS2: R1 v R2 |
|  | CS3-O3 | CS3: R1 v R2 |

#### Instrumental Training

Rats were next trained to press two separate levers to earn two different outcomes (R1-O1; R2-O2). Of the three pellets introduced during food cup training, two were used for these distinct R-O associations. Lever outcome assignments were counter-balanced across rats. At first, pellets were delivered on schedule of continual reinforcement (CRF), such that every press earned a single pellet. Rats were required to reach an acquisition criterion of earning 50 pellets within 40 min, before transitioning to variable interval (VI) schedules of reinforcement. Each lever was trained in isolation and rats were only transitioned to VI training once reaching the acquisition criteria on both levers. VI training sessions lasted for 45 min. For the first 20 min of these sessions one lever was available. This lever was then retracted and after a five min break during which neither lever was present, the other lever was inserted and remained available for the final 20 min of the session. During these sessions, VI reinforcement schedules were executed as follows: the first lever press occurring after passage of a pre-selected interval of time resulted in delivery of two pellets, triggering selection and initiation of a new interval. The VI schedules were increased slowly across 8 sessions of training in the following sequence: VI10 (range: 5-15sec), VI30 (range: 15-45sec), VI45 (range: 30-60sec), and VI60 (range: 45-60sec). Rats were trained for two sessions under each VI schedule. The first lever trained of the day was counterbalanced across session using a double alternating pattern (e.g., first lever trained of the day: L1, L2, L2, L1, L1, etc.).

#### Pavlovian Conditioning

Following instrumental training, rats were conditioned to associate three unique CSs with three different food pellet outcomes (CS1-O1; CS2-O2; CS3-O3). Importantly, CS1 and CS2 share a common outcome with the instrumental R1 and R2, therefore these associations were designed to capture SS PIT (see below). In contrast, CS3 is paired with O3, an outcome not shared by either lever and therefore this CS3-O3 association was designed to capture General PIT. All three CSs were auditory stimuli presented for 120 sec. During CS presentations, four food pellets were randomly delivered into the food cup on a VT20 schedule (range 11-30sec). This delivery schedule ensured that pellets were never delivered within the first 10 sec of CS presentation; this allowed us to measure anticipatory conditioned food cup approach without interference of consummatory behaviors. A white noise (60 dB), a tone (57 dB), and a click train (20 Hz) were each used as the CSs. Each CS was trained in isolated sessions that lasted 30 min and consisted of four CS-O trials separated by a variable five min inter-trial-interval (ITI; range: 3-7min). CS-O assignments were counterbalanced to ensure that each stimulus and flavor was evenly represented in SS and General CS-O associations within each group. Each session was separated by ∼40 min and rats underwent three separate sessions per day (one for each CS). Throughout Pavlovian conditioning, levers were unavailable and pellet delivery was not contingent upon any behavioral response. Food cup entries were recorded throughout.

#### Pavlovian to Instrumental Transfer Testing

PIT testing was conducted 2 and 4 days after the last Pavlovian conditioning session. Rats were given an instrumental *reminder* training identical to training sessions described above, the day prior to each PIT test. PIT testing lasted for 44 min, both levers were available throughout, but no pellets were delivered within the session. The session began with simultaneous insertion of both levers into the chambers. After 10 min, each of the three CSs was presented three times in a quasi-random order, with presentations separated by a fixed 2-min ITI. Lever presses, food cup entries and video footage were recorded throughout. Each rat was tested two times with one day of instrumental reminder training in between.

These training and testing procedures were designed to capture two distinct forms of PIT, SS and General PIT (e.g., Corbit and Balleine, 2005). SS PIT is observed when presentation of the sensory specific CSs (CS1 or CS2) results in greater responding on the lever that previously generated the same outcome versus the other lever that generated a different outcome than that predicted by the CS. Thus, the critical behavioral feature defining SS PIT is the differential influence of CS presentation on the rate of lever responding between the *Same* and *Different* levers. In contrast, General PIT is observed when presentation of a CS augments lever responding for an outcome not explicitly predicted by that CS (illustrated in Fig 2B). In the current experimental design, rats were explicitly trained with a General CS (CS3) that was paired with an outcome that was never paired with an instrumental response. This procedure was designed to maximize our ability to observe General PIT, by measuring the effect of CS3 presentations on lever responding. However, it is important to note that General PIT can also be observed during presentation of the sensory specific CSs (CS1 and CS2), as responding on the Different lever greater than pre-CS rates of responding. The 60 sec immediately preceding CS presentation was defined as the pre-CS period.

**FIG. 1:**
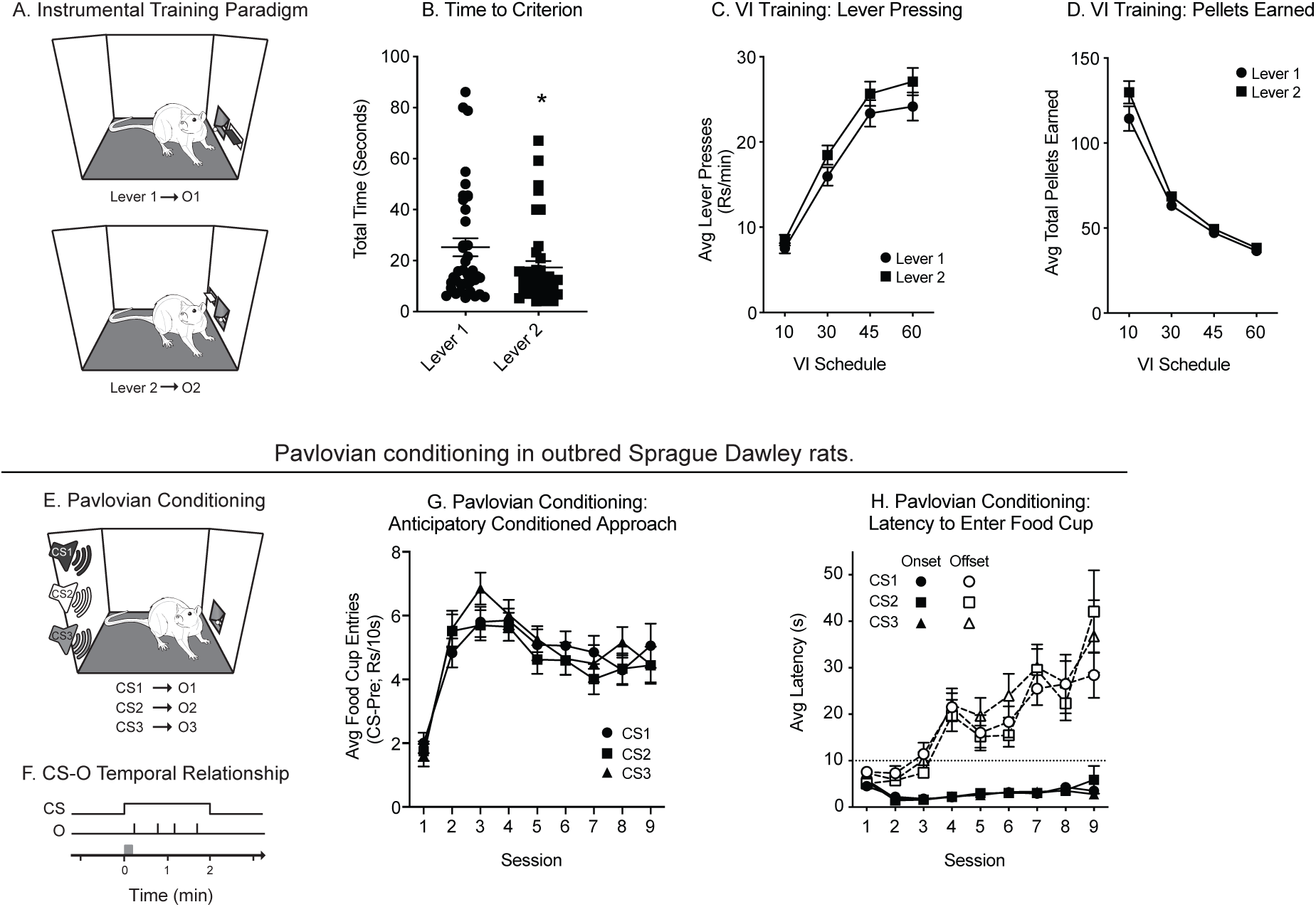
Instrumental and Pavlovian conditioning in outbred rats; N=39. **A)** Schematic of instrumental training. **B)** Total time to reach acquisition criterion during continual reinforcement training. **C)** The average rate of lever responding during variable interval (VI) instrumental training increased as the VI lengths increased. **D)** The average number of pellets earned decreased across VI training. **E)** Schematic of Pavlovian training. **F)** Schematic of the CS-Outcome relationships, depicting the delivery of four pellets within each 2-min CS presentation. The grey box over the timescale illustrates the 10-sec window during which no pellets are delivered following CS onset. **G)** Anticipatory conditioned approach during the first 10 seconds of CS presentation increased between session 1 and 2, remained stable thereafter, and was similar between CSs. **H)** The latency to enter the food cup following CS onset was rapid and stable across training, whereas the latency to enter following CS offset increased across training. All data are shown as averages ±SEM, unless otherwise noted.

**FIG. 2:**
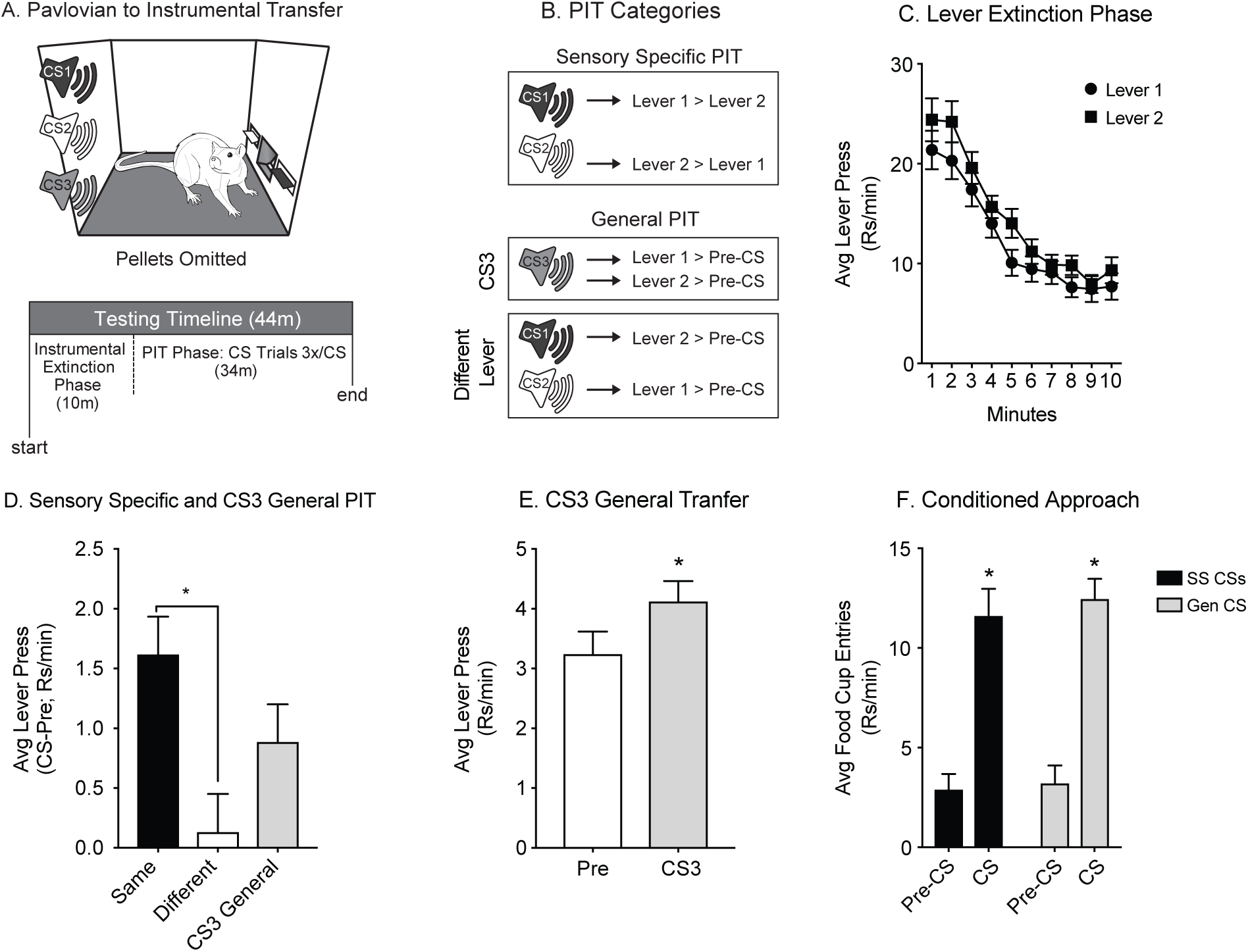
Expression of SS and General PIT in outbred rats. **A)** Schematic of PIT testing. **B)** Assessment of Sensory Specific PIT and General PIT. Sensory Specific PIT arises when a CS preferentially invigorates lever responding on the lever that shared an outcome with that CS. General PIT arises when responding for an outcome not predicted by a CS invigorates lever responding above pre-CS levels. This can be seen as CS3 General Transfer to either lever, or as transfer to the levers predicting a different outcome in response to CS1 or CS2 presentation. **C)** Extinction of lever responding during the 10 min pre CS-presentation phase is similar across both levers. **D)** Outbred rats exhibited SS PIT, with greater rates of lever pressing on the Same vs. Different Lever. **E)** Outbred rats show General PIT, with increases in the rate of lever pressing above pre-CS levels elicited by CS3. **F)** Conditioned approach to the food cup during PIT testing is similar between SS and General CSs (*=p<0.05).

### B. Experiments

#### Experiment 1

The goal of Experiment 1 was to determine whether obesity vulnerability was linked to pre-existing differences in the expression of sensory specific versus affective motivation (i.e., SS versus General PIT, respectively). To assess this, outbred male rats were trained and tested as described above, immediately following testing, rats were relieved from food restriction and allowed to consumed *ad libitum* standard lab chow for 3 days and then placed onto a moderately fatty palatable Junk-Food diet with *ad libitum* access. The purpose of this diet manipulation was to identify individual propensity to weight gain to determine obesity vulnerable individuals (as in Robinson et al., 2015). The Junk-Food (JF) diet was a mash consisting of Chips Ahoy! chocolate chip cookies (16% w/w; 260g), Frito Lays potato chips (5% w/w; 80g), Jif peanut butter (16% w/w; 260g), Nestle Nesquik chocolate powder (16% w/w; 260g), Test Diet, 5001 (25% w/w; 400g) and water (22% w/w; 355ml). Food intake (per cage) and body weight were recorded daily. In addition, nuclear magnetic resonance analyses (NMR) were conducted on a subset of rats prior to being placed on the JF diet and again following 5 weeks of JF consumption. These scans were performed by the Metabolism, Bariatric Surgery and Behavior Core at the University of Michigan. NMR data was gathered using a Bruker Minispec LF 90II device to measure lean mass, fat mass and body fluids. Body weight at the end of the 5 weeks of JF diet access were used for correlational analyses. As control, parallel correlational analyses were conducted comparing pre-training *ad libitum* weights.

#### Experiment 2

The goal of Experiment 2 was to determine whether SS and/or General PIT were enhanced in selectively obesity-prone than obesity-resistant rats, to help clarify our previous finding that obesity-prone rats exhibited enhanced single outcome PIT. To achieve this, we trained and tested selectively bred obesity-prone and obesity-resistant rats using identical procedures to those in Experiment 1. No post-training diet manipulation was used in this experiment because the identity of individual obesity susceptibility was known *a priori* via selective breeding (chacterized in, Levin et al., 1997; Vollbrecht et al., 2015).

### Statistical Analysis

Data was processed and organized with Microsoft Excel (Version 16.16.16) and statistical analyses were performed using the GraphPad statistical software suite Prism (Version 8.02). Data were analyzed using students t-tests, One-way, Two-way and Three-way repeated measures ANOVAs (RM ANOVAs) and Holms and Sidak’s multiple comparison tests for planned and post-hoc multiple comparisons. Correlational analyses were conducted using Pearson product-moment correlation coefficients.

Continual reinforcement training data were analyzed using the total time to reach the acquisition criteria per lever. The total time to acquire was summed across sessions for each lever. For instrumental responding, data were analyzed by obtaining average rates per session, and then averaging these rates across all sessions within each VI. Lever pressing data are presented as average responses per min (Rs/min) and pellets earned as averages per session. Behavior during Pavlovian conditioning was analyzed by obtaining session averages of anticipatory conditioned approach and latencies to approach the food cup following CS onset and offset. Anticipatory conditioned approach was evaluated by subtracting the number of food cup entries during the 10 sec pre-CS period from the first 10 sec of CS presentations (Rs/10s). Data from PIT testing were analyzed as responses per minute, with 60 sec pre-CS responding subtracted from CS responding when relevant and then averaged across trials, and tests.

## III. RESULTS

### A. Exp 1: Individual differences in outbred rats

#### Instrumental Training

Rats were trained to make an instrumental action to receive food pellets, with each of two separate levers earning one distinct type of food pellet outcome (Lever1-O1 and Lever2-O2; Fig 1A). Rats were trained under continual reinforcement conditions (CRF; one response leading to one pellet) until they reached the acquisition criterion (50 consecutive pellets in less than 40 min) for each lever. Despite full counter-balancing, rats reached the acquisition criterion slightly more quickly for Lever 2 than Lever 1 (Fig 1B: Paired t-test, Lever 1 vs. Lever 2, *t*_(37)_=3.16, p<0.01). During the next phase of instrumental training, rats were shifted to a variable interval (VI) schedule of reinforcement where the VI lengths were increased over 8 days of training. As expected this resulted in a steady increase in rates of responding as the VI lengths increased (Fig 1C: Two-way RM ANOVA: main effect of schedule: *F*_(3,114)_=126.3, p<0.01). The number of pellets earned across training decreased systematically as the VI length increased (Fig 1D: Two-way RM ANOVA: main effect of schedule: *F*_(3,114)_=317.1, p<0.01; no lever × schedule interaction, p=0.15). While we did observe slight differences in response rates between the levers, the outcome assignments and position of these levers was counterbalanced across rats; therefore, it is unlikely this effect reflects a meaningful difference between these responses (Fig 1C: Two-way RM ANOVA: main effect of lever: *F*_(1,38)_=6.75, p=0.01; no schedule × lever interaction, p=0.20). Moreover, the number of pellets earned between the levers was similar across training (Fig 1D: Two-way RM ANOVA: no main effect of lever, p=0.08; Holms and Sidak’s (HS) multiple comparisons, VI60: Lever 1 v Lever 2, p=0.87). Together these data demonstrate stable acquisition of instrumental lever responding across VI training for two distinct response-outcome associations.

#### Pavlovian Conditioning

After instrumental training, rats were conditioned with three distinct CS-outcome relationships (Fig 1E). On each trial, the CS was presented for 120 secs during which four pellets were delivered into the food cup, with pellet delivery never occurring within the first 10 sec of the CS (Fig 1F; grey box). This enabled us to examine CS-driven, anticipatory approach during the first 10 sec of CS presentation, prior to pellet delivery. This anticipatory approach behavior rapidly increased between first two sessions and then stabilized for the remaining sessions, and was similar across CSs (Fig 1G: Two-way RM ANOVA: main effect of session: *F*_(8,304)_=18.95, p<0.01; no effect of CS: p=0.42; no session × CS interaction, p=40). As an additional measure of conditioning, we also examined the latencies to approach the food cup following CS onset and offset. While initially the approach latencies between CS onset and offset were comparable, across conditioning sessions the approach latencies following CS offset slowed dramatically (Fig 1H open symbols: Two-way RM ANOVA: phase × session interaction, CS1: *F*_(8,304)_=5.18, p<0.01; CS2: *F*_(8,304)_=11.79, p<0.01; CS3: *F*_(8,304)_=7.73, p<0.01). In addition, the latency to approach the food cup following CS onset was significantly faster than approach following CS offset (Fig 1H: Two-way RM ANOVA: main effect of phase, CS1: *F*_(1,38)_=92.88, p<0.01; CS2: *F*_(1,38)_=137.4, p<0.01; CS3: *F*_(1,38)_=67.19, p<0.01). Collectively, these data show that acquisition of anticipatory conditioned approach is similar across three distinct CS-outcome associations.

#### Pavlovian to Instrumental Transfer Testing

Following instrumental and Pavlovian conditioning, rats were tested for PIT under extinction conditions. Each test session began with simultaneous insertion of both levers, which remained available throughout testing. After an initial instrumental extinction phase (10 min), rats were presented with each CS three times, delivered in a quasi-random order (Fig 2A). Lever responding steadily declined across the first 10 min of testing, as expected, and rates of responding were similar between both levers across time (Fig 2C: Two-way RM ANOVA: main effect of time: F_(9,342)_=91.84, p<0.01; no effect of lever, p=0.07; no lever × time interaction, p=0.97). Classically, SS PIT is observed when presentation of a CS results in greater responding on the lever that previously delivered the same outcome than on the lever that previously delivered a different outcome than that predicted by the CS (i.e., Same Lever ¿Different Lever). This effect is captured by presentation of CS1 and CS2 (Fig 2B). In contrast, General PIT is observed when presentation of a CS increases lever pressing on a lever that previously did not share an outcome with that predicted by the CS. This effect is classically captured by responding above baseline (pre-CS) in response to presentation of the CS3 (which does not share an outcome with either lever, CS3 General PIT), but can also be measured by lever pressing on the Different Lever in response to CS presentation (Fig 2B; Different Lever General PIT).

Analyses of lever response data revealed that rats showed substantial SS PIT, preferential responding on the lever that produced the same outcome than on the lever that produced a different outcome to that predicted by the CS being presented (Fig 2D: One-way RM ANOVA: main effect of time: *F*_(1.62,61.54)_=5.67, p<0.01; HS multiple comparisons, Same vs. Different, *t*_(38)_=2.83, p=0.02). In addition, the magnitude of General transfer in response to CS3 presentation was not significantly different from that of Same and Different transfer (Fig 2D: HS multiple comparisons, Same vs. General, *t*_(38)_=1.65, p=0.11; Different vs. General, *t*_(38)_=2.25, p=0.06). Rats also showed General PIT, as indicated by a significant increase in the rate of lever responding during CS3 presentation above pre-CS responding (Fig 2E: Paired t test, Pre v CS, *t*_(39)_=2.78, p<0.01). Together these data confirm the expression of both SS and General PIT following the training and testing conditions used here (See Fig 3 for data related to Different Lever PIT).

**FIG. 3:**
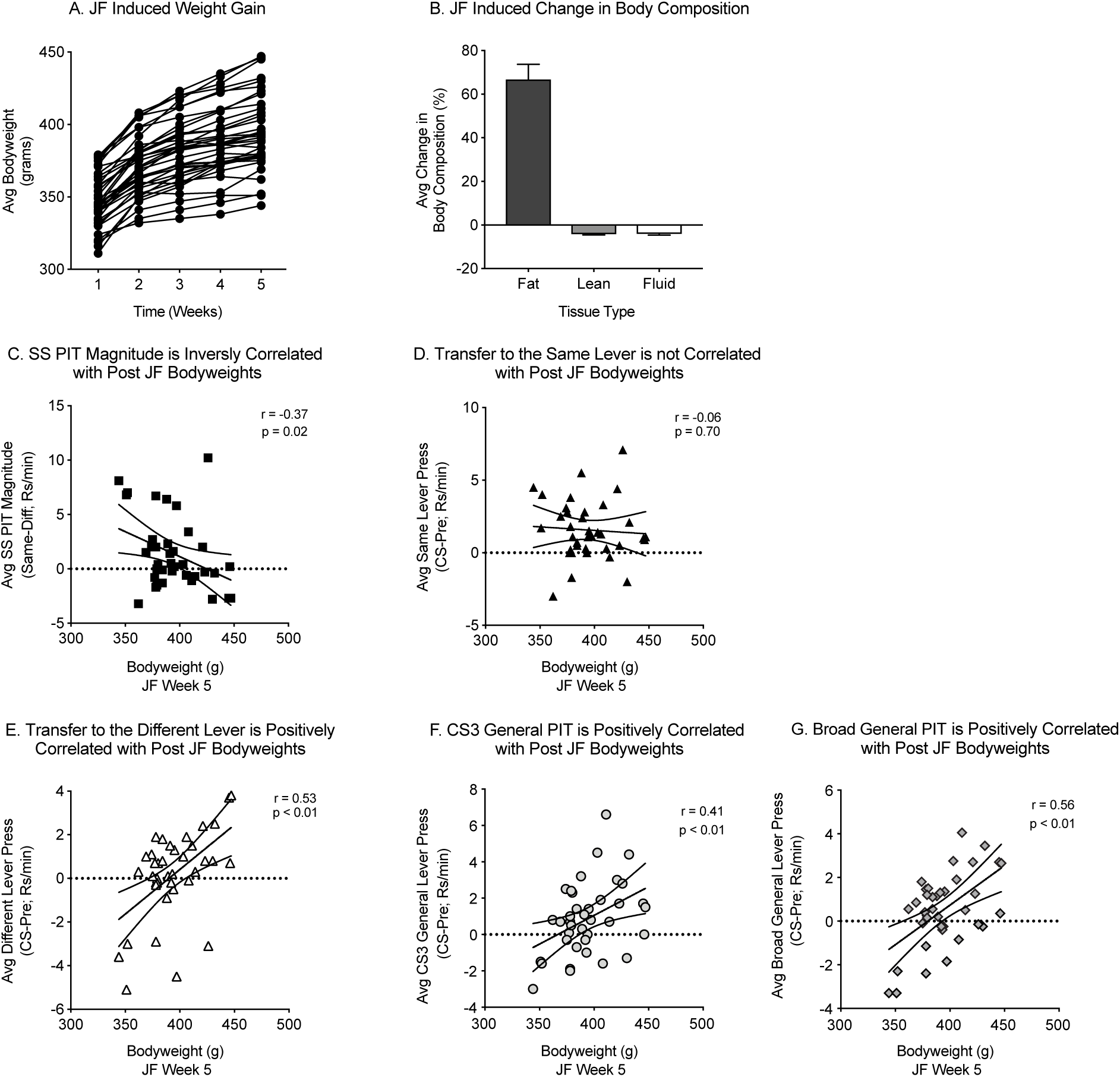
Post junk-food weight is positively correlated with the magnitude of General PIT, but not SS PIT. **A)** Junk-food diet consumption produced a range of weight gain across the five week diet exposure. **B)** Average percent change in fat, lean, and fluid mass from baseline following junk-food consumption. **C)** SS PIT magnitude (i.e., the difference between transfer to the Same vs. Different levers) is negatively correlated with weight. **D)** Transfer to the Same lever did not correlate with post junk-food weight. **E)** Transfer to the Different Lever was strongly correlated with post junk-food weight. **F)** CS3 elicited General Transfer was strongly correlated with subsequent weight gain. **G)** Broad General PIT (CS3 and Different transfer) was also strongly correlated with post junk-food weight.

In addition to measuring PIT, we also recorded food cup entries during PIT testing as a measure of conditioned approach. To provide an indication of whether response competition between food cup entries and lever responding differs between CS types (SS: CS1 and CS2; General: CS3), conditioned approach data were analyzed according to CS type (Fig 2F). Although SS and General CSs supported different rates of lever responding (Fig 2D), these CSs supported similar magnitudes of conditioned approach to the food cup (Fig 2F: Two-way RM ANOVA: main effect of phase: *F*_(1,15)_=151, p<0.01; no effect of CS type, p=0.54; no phase × CS type interaction, p=0.74). Thus, there does not appear to be substantial response competition between food cup entries and lever responding or across SS and General CSs.

#### Post Training Junk-Food Consumption and Weight Gain

We next examined the relationship between the expression of PIT and subsequent weight following free access to junk-food diet (see methods for details).All rats gained a significant amount of weight across the 5 weeks of junk-food consumption (Fig 3A: One-way RM ANOVA: main effect of time: F_(1.35,49.99)_=252, p<0.01). In final week of *ad libitum* junk-food access, the average weight was 394g (SEM: 4.2g) with a range of 103g. For a subset of rats, body composition was determined before and after junk-food consumption. As expected, junk-food consumption produced a significant increase in body fat mass, a decrease in lean mass and had no effect on fluid mass compared to baseline (Two-way RM ANOVA: main effect of baseline vs. post: F_(1,18)_=15.20, p<0.01; main effect of mass type: F_(1.04,18.69)_=30504, p<0.01; timepoint × mass type interaction, F_(1.04,18.68)_=103.5, p<0.01). These data are summarized in Fig 3B as the average percent change from baseline. Collectively, these data demonstrate that junk-food induced weight gain and increased fat mass, with a wide range in the magnitude of in the final weights reached

#### Relationships between PIT Magnitude and subsequent weight following junk-food consumption

We next examined relationships between weight following junk-food consumption and the magnitude of SS- and General PIT. SS PIT magnitude (i.e., the difference in CS elicited responding on the Same vs. Different levers) was significantly inversely correlated with subsequent weight gain (Fig 3C: Pearson correlational analysis: r=−0.37, p=0.02, n=38). However, given that SS PIT magnitude is comprised of two variables, Same transfer and Different transfer, we also directly examined correlations between weight and response rates on the Same Lever or the Different Lever in separate analyses. This revealed that response rates on the Same Lever were not significantly correlated with post junk-food weights (Fig 3D: Pearson correlational analysis: r=−0.06, p=0.70, n=38), whereas there was a strong positive correlation between weight and response rates on the Different Lever (Fig 3E: Pearson correlational analysis: r=0.53, p<0.01, n=38). Thus, the initial negative correlation when considering the SS PIT difference score arises because subtraction of a positive relationship (Different transfer) from the absence of any relationship (Same transfer) results in the appearance of a negative correlation. This also suggests that differences in General PIT magnitude may be more tightly linked to obesity susceptibility, because transfer to the Different Lever is a form of General PIT (see below and discussion). Consistent with this, CS3-elicited General PIT was strongly positively correlated with post junk-food weight (Fig 3F: Pearson correlational analysis: r=0.41, p<0.01, n=38).

General PIT is classically defined as the ability for a CS to augment instrumental responding for an outcome that is not directly predicted by the CS presented. The two critical features in this operational definition are that 1) instrumental responding must be increased above baseline levels and that 2) the instrumental response must be for an outcome other than that predicted by the CS. Thus, in addition to CS3 elicited responding, General PIT is also captured when presentation of CS1 or CS2 elevates responding on the Different Lever. In defense of this concept, we found a strong positive correlation between Different Lever transfer and transfer during CS3 presentations (Data not shown: Pearson correlational analysis: r=0.45, p<0.01, n=38). Therefore, we collapsed across CS3 elicited and Different Lever responding to obtain a more complete measure of General PIT (i.e., Broad General PIT). This revealed an even stronger correlation between Broad General PIT and weight compared to relationships to Different Lever or CS3 responding alone (Fig 3G: Pearson correlational analysis: r=0.56, p<0.01, n=38). Critically, the same analyses performed using pre-training weights were not correlated with PIT magnitude (Data not shown: Pearson correlational analysis: SS PIT magnitude, r=0.04, p=0.80, n=38; Same transfer, r=0.05, p=0.73, n=38; Different transfer, r<0.01, p=0.96, n=38; CS3 General transfer, r=0.06, p=0.71, n=38; CS3 Broad General transfer, r=0.03, p=0.86, n=38). Thus, it is not simply that rats that were heaviest at the time of PIT testing show the strongest magnitude of PIT, but rather that susceptible rats show stronger General PIT magnitude prior to diet exposure.

### B. Exp 2: Expression of SS and General PIT in selectively bred obesity-prone and obesity-resistant rats

#### Instrumental Training

As in experiment 1, following food cup training, rats were trained to press one lever to earn one outcome and another lever to earn a different outcome using a CRF schedule with a criterion of earning 50 consecutive outcomes in under 40 min. Obesity-resistant and obesity-prone rats reached this criterion in a similar amount of time on both levers (Fig 4A: Two-way RM ANOVA: no effect of group, p=0.57; no effect of lever, p=0.20; no group × lever interaction, p=0.33). The average time to reach the acquisition criterion on both levers was min (SEM: ±4.37). Rats then began training under VI schedules. During this phase, the rate of lever pressing increased as the VI duration was increased across training, and rates of responding did not differ between levers (Fig 4B: Three-way RM ANOVA: main effect of VI schedule, *F*_(3,84)_=80.92, p<0.01; no effect of lever, p=0.65; no lever × schedule interaction, p=0.14). In addition, across VI training rates of lever responding in obesity-prone rats increased relative to obesity-resistant rats (Fig 4B: Three-way RM ANOVA: main effect of group, *F*_(1,28)_=4.78, p=0.03; group × schedule interaction, *F*_(3,84)_=3.09, p=0.03; no schedule × group × lever interaction, p=0.17). The number of pellets earned decreased across the VI schedules, as expected (Fig 4C: Three-way RM ANOVA: main effect of schedule, *F*_(3,84)_=69.88, p<0.01). However, despite group differences in lever response rates, both obesity-resistant and obesity-prone groups earned similar amounts of pellets across VI training (Fig 4C: Three-way RM ANOVA: no effect of group, p=0.97; no effect of lever, p=0.96; no group × lever interaction, p=0.41; no group × lever × schedule interaction, p=0.12). In sum, both obesity-prone and obesity-resistant rats similarly acquire instrumental lever responding on two separate levers and, although there were statistical differences in lever response rates during VI training (mean difference VI60: 6.35, SEM: 3.03), both groups earned a similar number of pellets across training sessions.

**FIG. 4:**
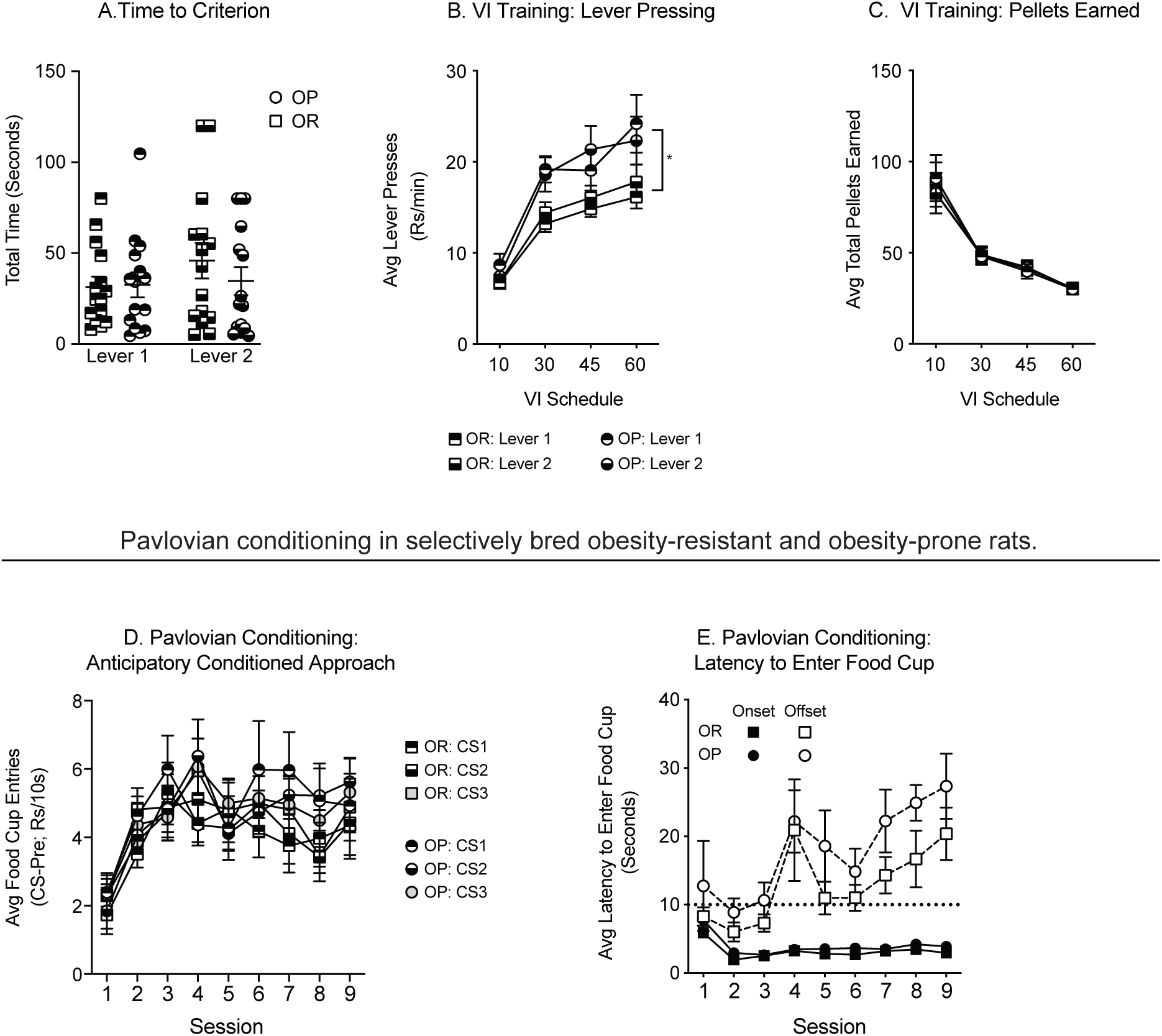
Instrumental and Pavlovian conditioning in selectively bred rats (OP N=15, OR N=15). **A)** Total time to reach acquisition criteria during continual reinforcement training was similar between groups. **B)** The average rate of lever responding during variable interval (VI) instrumental training increased as VI lengths increase, and is greater in obesity-prone vs. obesity-resistant rats. **C)** The average number of pellets earned decreased across VI training and was similar between groups. **D)** Anticipatory conditioned approach during the first 10 seconds of CS presentation increased between sessions 1-3 and remained stable thereafter in both groups. Approach was similar between groups and CSs. **E)** The latency to enter the food cup following CS onset became faster between sessions 1 and 2, was stable across the remaining sessions, and did not differ between groups. Latency to enter the food cup following CS offset slowed across sessions and did not differ between groups (*=p<0.05).

#### Pavlovian Conditioning

Following instrumental training, rats were conditioned in 9 sessions to associate three distinct CSs with distinct outcomes, as above. Anticipatory conditioned approach within the first 10 seconds of CS presentation increased across training in both groups (Fig 4D: Two-way RM ANOVA: main effect of session, OP: *F*_(8,112)_=5.55, p<0.01; OR: *F*_(8,112)_=3.70, p<0.01) and did not differ across CSs in either group (Fig 4D: Two-way RM ANOVA: no effect of CS, OP: p=0.96; OR: p=0.68; no CS × session interaction, OP: p=0.47; OR, p=0.48). Additionally, rates of food cup entries were similar between obesity-prone and obesity-resistant groups (Fig 4D: Two-way RM ANOVA: no effect of group, CS1: p=0.28; CS2: p=0.62; CS3: p=0.64; no group × session interaction, CS1: p=0.63; CS2: p=0.27; CS3: p=0.99). We also examined latencies to enter the food cup following CS onset and offset. For ease of presentation, these data are collapsed across CSs given that no CS effects were observed (data not shown: Two-way RM ANOVA: no effect of CS, Onset: OP: p=0.47; OR: p=0.77; Offset: OP: p=0.80; OR: p=0.11). As expected, rats displayed rapid food cup approach following CS onset, and as conditioning progressed approach following CS offset slowed (Fig 4E open symbols: Three-way RM ANOVA: main effect of session, *F*_(8,224)_=5.58, p<0.01; main effect of phase, *F*_(1,28)_=69.98, p<0.01; session × phase interaction, *F*_(8,224)_=6.46, p<0.01). Furthermore, latencies were similar in obesity-prone and obesity-resistant rats (Fig 4E: Three-way RM ANOVA: no effect of group, p=0.07; no group × phase interaction, p=13; no group × phase × session interaction, p=0.97). These data confirm that obesity-prone and obesity-resistant rats acquired the CS-outcome associations without any notable differences in acquisition between the groups.

#### Pavlovian to Instrumental Transfer Testing

Rats were tested for PIT following training using procedures identical to Experiment 1. During the first 10 min of testing, lever responding declined as expected, and lever response rates did not differ between groups (Fig 5A: Three-way RM ANOVA: main effect of time, *F*_(9,252)_=7,12, p<0.01; no effect of group, p=0.41). In addition, despite differences during VI training, rates of responding on both levers were similar between groups (Fig 5A: Three-way RM ANOVA: no lever × group or lever × group × time interaction). Next, CSs were presented and SS PIT and General PIT were assessed as described in Experiment 1. CS presentation evoked significantly more lever responding in obesity-prone rats than in obesity-resistant rats regardless of transfer type (Fig 5B: Two-way RM ANOVA: main effect of group, *F*_(1,28)_=8.33, p<0.01; no group × transfer interaction, p=0.23). In this experiment, we found substantial general transfer to the Different Lever which masked SS PIT (Holland, 2004). Specifically, while transfer to the Same Lever was substantially greater than CS3-elicited General transfer, Same Lever responding did not significantly differ from transfer expressed by responding on the Different Lever, the classic definition of SS PIT (Fig 5B: Two-way RM ANOVA: main effect of transfer, *F*_(2,56)_=3.86, p=0.03; HS multiple comparisons: Same vs. Diff, p=0.21). Thus, SS PIT as classically defined by the difference in Same and Different transfer was not apparent in this experiment. Given that Broad General PIT was most tightly correlated to obesity susceptibility in outbred rats, we next examined differences in Broad General PIT between selectively bred groups. We found that obesity-prone rats exhibited significantly greater Broad General PIT than obesity-resistant rats (Fig 5C: Two-way RM ANOVA: main effect of phase, *F*_(1,28)_=94.79, p<0.01; main effect of group, *F*_(1,28)_=6.41, p=0.01; phase × group interaction, *F*_(1,28)_=5,45, p=0.02; HS multiple comparisons: Pre v CS: OR *t*_(28)_=5.23, p<0.01; OP *t*_(28)_=8.53, p<0.01; OR v OP: Pre-CS, p=0.13; CS: *t*_(56)_=3.19, p<0.01).

**FIG. 5:**
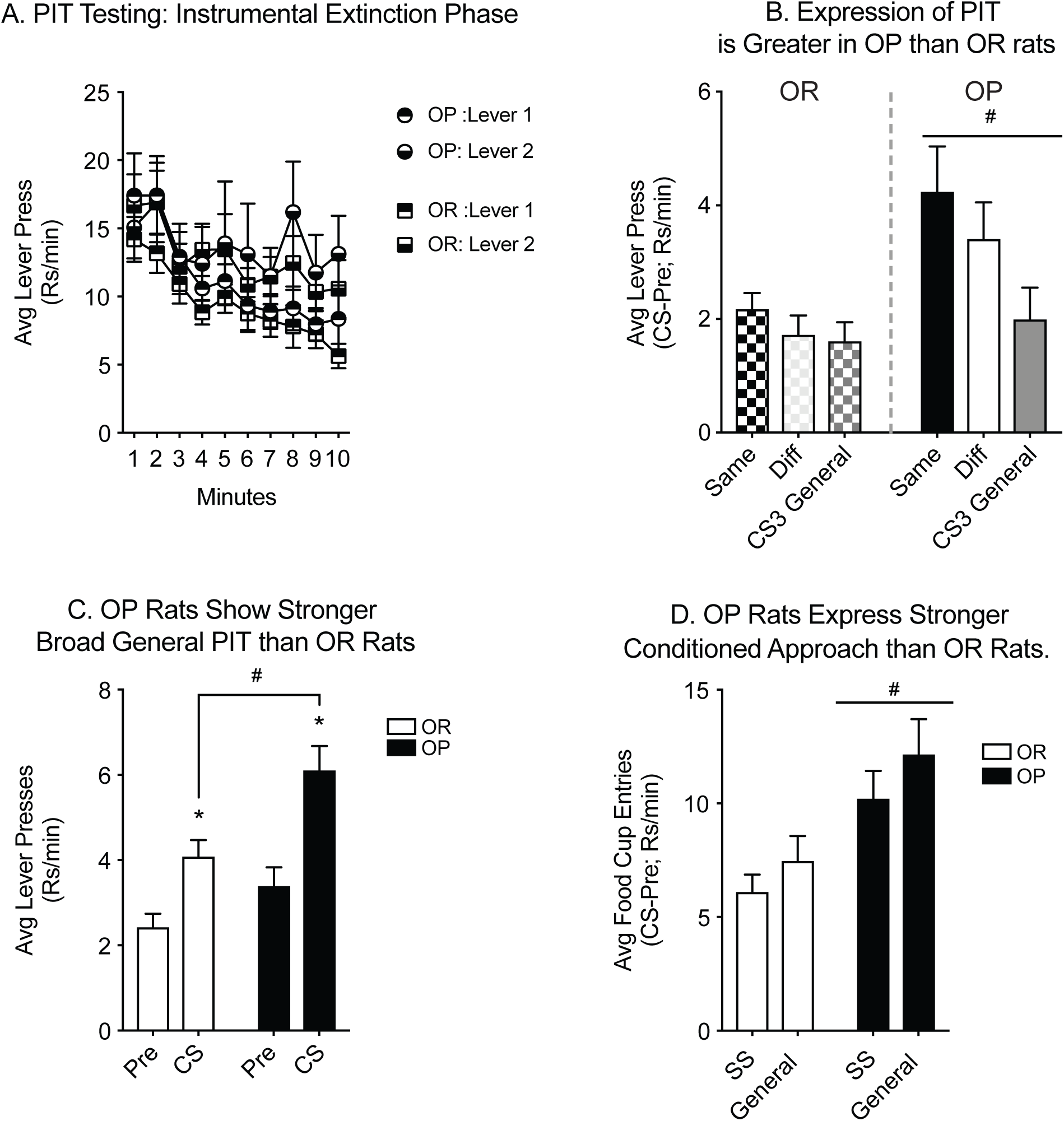
The magnitude of PIT and conditioned approach are greater in obesity-prone vs. obesity-resistant rats **A)** Lever pressing decreased in the first 10 minutes of testing, prior to CS presentation, and did not differ between groups. **B)** The magnitude of PIT was greater in obesity-prone rats, although both groups showed substantial General transfer to the Different Lever, masking Sensory Specific PIT. **C)** Both groups exhibited Broad General PIT, but this effect was stronger in obesity-prone rats. **D)** Conditioned approach was greater in obesity-prone rats, and did not differ between SS and General CSs in either group (*=p<0.05; #=p<0.05).

## IV. DISCUSSION

We previously found that selectively bred obesity-prone rats show stronger PIT than obesity-resistant rats prior to the onset of obesity (Derman and Ferrario, 2018). However, whether differences were driven by a sensory specific or general affective processes were unknown, and results had not been verified across models of individual susceptibility to obesity. In the current study, we asked whether susceptibility to obesity was associated with the propensity to exhibit Sensory Specific and/or General PIT using both outbred and selectively-bred rodent models. In outbred rats, we found that General PIT prior to weight gain was positively correlated with subsequent bodyweight following 5 weeks on a junk-food diet. Consistent with these data, we also found that selectively bred obesity-prone rats exhibited greater General PIT than obesity-resistance rats, prior to obesity.

### A. Obesity Vulnerability in Outbred Rats is Associated with Enhanced Affective Motivation

Obesity vulnerability is associated with enhanced brain and behavioral responsivity to food related stimuli, the latter of which may be one of the key behaviors that renders this vulnerability (Boswell and Kober, 2016; Stice et al., 2013). However, the process by which Pavlovian stimuli come to exert control over behaviors, particularly instrumental behaviors, can arise via at least two distinct mechanisms, a sensory specific process and a general affective process (Corbit and Balleine, 2005; Corbit and Balleine, 2011; Corbit et al., 2007). In Experiment 1, we determined the relationship between the magnitudes of SS and General PIT and subsequent weight following a junk-food diet. Rats were first taught two instrumental associations and then, in a separate phase, three Pavlovian associations (see Table 1). Following this training, they were tested for SS and General PIT by presenting the Pavlovian CSs in the presence of both levers (under extinction conditions). As expected, rats displayed both SS and General PIT (Fig 2D-E). To our knowledge, this is the first time this procedure has been used successfully outside the laboratory of the group that originally developed it, though some modifications to the original training procedure were made (see methods).

Following testing, rats were placed on junk-food diet for 35 days. This moderately fatty diet (19.6% fat) was designed to allow us to detect individual differences in weight gain, as diets with higher fat content (40-60%) tend to induce robust obesity in the majority of subjects. In addition, we have previously found that 30 days of exposure to this diet is needed to reliably identify individual differences (Oginsky et al., 2016a; Robinson et al., 2015). Bodyweight at the end of this diet manipulation was then used to assess the relationships between initial SS PIT and General PIT magnitudes and obesity susceptibility. Classically, SS PIT is defined by the differential influence of CS presentation on instrumental responding for the Same versus a Different outcome than that predicted by the CS (Same > Diff; Colwill and Motzkin, 1994). General PIT, on the other hand, is more broadly defined as an increase in instrumental responding for an outcome not specifically associated with the CS that is being presented. Thus, in our procedure, responding on either lever during presentation of CS3 (which does not share an outcome with either lever) or responding on the Different Lever (i.e., CS1-Lever2 or CS2-Lever1) both capture General PIT (here termed Broad General PIT). Consistent with this, there is a significant positive correlation between responding on the Different Lever and CS3-elicited lever responding (see results), substantiating the use of CS3 and Different Lever responding as a measure of Broad General PIT.

Examination of relationships between weight and SS PIT revealed a significant negative correlation between SS PIT magnitude and weight (Fig 3C). However, when relationships between weight and responding on the Same and the Different levers were examined separately, we found that transfer to the Same Lever was not correlated with weight (Fig 3D), but that transfer to the Different Lever was strongly positively correlated to weight (Fig 3D-E). Thus, the initial negative correlation of SS PIT magnitude arose because subtraction of a positive relationship (Different responding) from the absence of any relationship (Same responding) resulted in the appearance of a negative correlation. This also indicates that the main driver of these relationships was responding on the Different Lever. In terms of behavioral interpretation, these data suggest that obesity susceptibility is associated with stronger general affective incentive motivation, but is not strongly related to sensory specific incentive motivation. Consistent with this, there was also a strong positive correlation between weight and General PIT as measured by CS3 evoked responding (Fig 3F). When we collapsed across CS3-elicited lever responding and Different Lever responding to attain one metric of Broad General PIT, this positive relationship between weight and general PIT magnitude became even stronger (Fig 3G). Together these findings indicate that susceptibility to obesity is accompanied by enhanced affective motivation in outbred rats.

### B. Selectively Bred Obesity-Prone Rats Exhibit Enhanced Affective Motivation

While studies in outbred populations are valuable, the use of this outbred model to study basal mechanisms or pre-existing neuronal differences that render individuals vulnerable can be exceedingly challenging, and in some cases impossible. Fortunately, the development of selectively bred lines enables this type of research by identifying *a priori* individuals who are susceptible versus resistant (Alonso-Caraballo et al., 2018). Previous work from our lab has demonstrated that obesity-prone rats exhibit stronger single outcome PIT than obesity-resistant rats (Derman and Ferrario, 2018). However, this variant of PIT does not distinguish between sensory specific and affective mechanisms of control, hence the underlying psychological mechanism of this effect remained unclear. Thus, in Experiment 2 we determined the degree to which enhancements in affective Pavlovian motivation are found in selectively bred obesity-prone and obesity-resistant rats. In addition, this provides cross-model corroboration of phenotypic differences in obesity susceptible versus resistant populations.

Rats were trained and tested identically to the outbred rats from Experiment 1. During PIT testing, obesity-prone rats exhibited much stronger transfer effects than obesity-resistant rats on all three transfer measures (Same, Different, and CS3 General; Fig 5B). Moreover, Broad General PIT was stronger in obesity-prone versus -resistant rats (Fig 5C). Thus, stronger affective Pavlovian motivation appears to be the primary driver of differences between obesity susceptible versus resistant populations. These data are consistent with stronger single outcome PIT in obesity-prone versus obesity-resistant rats (Derman and Ferrario, 2018), and with results in outbred rats discussed. Jointly, these experiments identify enhanced Pavlovian affective motivation as a common mediator of stronger cue-triggered food-seeking in obesity susceptible populations.

One outstanding consideration of the PIT data in Experiment 2 is that the presence of robust General transfer to the Different Lever masked our ability to measure SS PIT by the classically defined comparison between Same and Different responding. Thus, data here do not rule out the possibility that there may be circumstances under which SS PIT differs between obesity-prone and obesity-resistant groups. This could be addressed by using procedures that may facilitate the expression of SS PIT over General transfer to the Different Lever. For example, in the current study CSs were paired with one of three differently flavored food pellets, where flavor and scent were the primary distinguishing sensory properties. However the use of more distinct outcomes, for instance liquid versus pellet reinforcers, is likely to promote sensory specific encoding and better capture SS PIT. Procedural modifications such as this may maximize the ability to observe differences in SS PIT between groups or following the onset of obesity in future studies.

Collectively, the data from Experiment 1 and 2 demonstrate that obesity susceptibility is associated with stronger General PIT. Thus, both naturally occurring susceptibility and susceptibility that is amplified through selective breeding are both accompanied by enhanced general affective Pavlovian motivation. Furthermore, neither outbred nor selectively bred rats exhibited notable enhancements in SS PIT, suggesting that the primary driver for enhanced Pavlovian incentive motivation in susceptible population arises particularly via a general affective process.

### C. Differences in Conditioned Approach Behavior

One noteworthy difference in the results from outbred versus selectivity bred rats here was that selectively bred obesity-prone rats showed stronger conditioned approach (Fig 5D) during PIT testing, whereas we did not observe any correlations between obesity vulnerability and the magnitude of conditioned approach in outbred rats. This discrepancy suggests that selective breeding for obesity susceptibility may have magnified the intensity of Pavlovian motivational control above and beyond that seen in obesity vulnerable outbred rats. In addition, it is worthwhile to note that in our previous single outcome PIT study, we did not find significant differences in conditioned approach between selectively bred obesity-prone and obesity-resistant groups during PIT testing (Derman and Ferrario, 2018). It is likely that this discrepancy arose from differences in the training paradigms between this and the current study. Notably, to assess single outcome PIT rats were conditioned using a Pavlovian discrimination task, where a CS+ was paired with pellets contrasted with a CS-that was never paired with pellets. In contrast, in the current procedure rats were trained with 3 distinct CSs, each paired with pellets. These conditioning paradigms differ by two relevant aspects. Discrimination conditioning entails some degree of inhibitory learning as rats learn to withhold conditioned approach to CS-presentations. The engagement of inhibitory processes in this procedure may have dampened the expression of enhanced conditioned approach in obesity-prone rats. Another related distinguishing feature between the current and previous study is that the density of outcomes within the conditioning session was significantly leaner in the previous study, where in the previous study, in a 60-minute session rats were presented with 16 total pellets across four CS+ trials. In contrast, in the current study 16 total pellets were presented in each 30-minute session and rats underwent 90 minutes of training in 3 sessions per day. Consequently, the total number of rewards experienced and the density of outcomes per session was much richer in the current experiment. It is likely that the richness of training in the current experiment enhanced the attribution of incentive salience to the CSs, and that this was most pronounced in obesity-prone rats due to their sensitivity to Pavlovian motivation. This finding is particularly interesting to consider in terms of ecological relevance of these effects, because it suggests that in environment replete with rewarding food experiences Pavlovian stimuli can exert differential motivational effects in vulnerable versus resistant individuals.

### D. Implications for Neurobiological Differences

As mentioned in the introduction, SS PIT and General PIT are mediated by distinct psychological and neurobiological processes. Thus, the stronger General PIT found in obesity vulnerable individuals points not only to stronger Pavlovian affective motivation (discussed above), but also to potential neurobiological differences between susceptible and resistant populations. Although not mechanistically well defined, lesion and inactivation studies have revealed that General PIT is mediated by the ventral tegmental area (VTA), the central nucleus of the amygdala (CN), and the NAc Core (Corbit and Balleine, 2005; Corbit and Balleine, 2011; Corbit et al., 2007). Thus, basal differences within these nuclei, or their connectivity, may exist between obesity susceptible and non-susceptible individuals. Additionally, there could also be differences in plasticity within these circuits induced by Pavlovian and instrumental training. Consistent with this possibility, Pavlovian and instrumental training for single outcome PIT are sufficient to increase the expression of calcium permeable AMPA receptors (CP-AMPARs) within the NAc of obesity-prone but not obesity-resistant rats. Furthermore, blockade of CP-AMPARs within the NAc core is sufficent to block the expression of single outcome PIT in obesity-prone populations (Derman and Ferrario, 2018). Finally, excitability (that is how readily neurons fire) is enhanced in obesity-prone versus obesity-resistant rats (Oginsky et al., 2016b), suggesting that the threshold for plasticity may be lower in obesity susceptilbe populations. Thus, together, these data suggest that similar experiences may promote distinct plasticity across neural circuits and/or within specific regions that mediate affective motivation in susceptible versus resistant populations.

In addition to the NAc core, the expression of General PIT also relies on activity within the CN of the amygdala, although it should be noted that there is no evidence for a direct connection between these brain regions. The CN is a GABAergic structure that receives glutamatergic inputs from several regions including the insular cortex, hypothalamus, midbrain, pons, and medulla; (Sah et al., 2003). Consequently, differences in experience-induced glutamatergic plasticity within the CN may promote enhanced incentive motivation as has been observed in the NAc Core (Conrad et al., 2008; Lee et al., 2013; Loweth et al., 2014; Ma et al., 2014; Wolf, 2016). In addition, in contrast to other amygdalar regions like the BLA, the CN receives dense dopamine inputs, and antagonism of D1 receptors in the CN potentiates cocaine self-administration (Freedman and Cassell, 1994; McGregor and Roberts, 1993). Thus, alterations in dopaminergic transmission in the CN may also play a role in mediating enhanced affective incentive motivation observed in susceptible individuals. Consistent with this idea, sensitization to cocaine-induced locomotion is enhanced in obesity-prone versus obesity-resistant rats (Oginsky et al., 2016b). This suggests that dopamine (DA) transmission may be enhanced more dramatically by experiences with reward in obesity-prone versus obesity-resistant groups. However, whether there are differences in glutamatergic or dopaminergic transmission within the CN that may contribute to behavioral differences observed here remains and open question.

## Summary and Broader Implications

In sum, the behavioral data using two different models of individual susceptibility to obesity show that affective Pavlovian motivation is enhanced in obesity susceptible compared to resistant populations. This, in conjunction with previous studies, identify enhanced incentive motivational responses to food cues as phenotypic difference in obesity susceptible versus resistant populations. In addition, these results point to potential differences in plasticity of amygdalar and striatal circuits that mediate PIT in these populations. Furthermore, these data may help explain why maintained weight loss and prevention of weight gain is particularly difficult for vulnerable individuals within the cue rich modern food environment. However, one unique feature of General PIT is that it is a particularly labile form of transfer that is sensitive to shifts in appetite, such that satiation abolishes General PIT (Balleine, 1994; Corbit et al., 2007). This suggests that prevention plans centered on maintaining the sensation of satiation (or helping to restore that sensation in obese individuals) may help to curb expression of enhanced incentive motivation in vulnerable populations. Finally, data here are consistent with human studies showing that that susceptibility to weight gain is associated with enhanced activity in the amygdala and NAc elicited by food cues (Demos et al., 2012; Yokum et al., 2014), as well as with the finding that obese and over-weight individuals show enhanced motivational responses to food cues, and greater cue-induced consumption (Rogers and Hill, 1989; Rothemund et al. 2007; Stice et al. 2008; Stoeckel et al., 2008; Bruce et al. 2010; Martin et al. 2010; Ng et al. 2011). Thus, results here provide unique insights into the underlying psychological and neurobiological mechanisms that drive food-seeking and overeating in susceptible individuals.

## Acknowledgments

This work was supported by the National Institutes of Health [NIDDK 1F31-DK111194-01 awarded to RCD and R01DK106188, R01DK115526 awarded to CRF].

## Author Contributions

RCD designed conducted experiments, analyzed data, and wrote the manuscript. CRF designed experiments, analyzed data, and wrote the manuscript.

